# A SNP panel for co-analysis of capture and shotgun ancient DNA data

**DOI:** 10.1101/2025.07.30.667733

**Authors:** Romain Fournier, Alice Pearson Fulton, David Reich

## Abstract

Advances in technology have decreased the cost of generating genetic variation data from ancient people, resulting in exponentially increasing numbers of individuals with whole genome data. However, each technology comes with platform-specific biases, limiting co-analyzability of individuals sequenced with different technologies as well as joint analysis of modern and ancient individuals. We present a method to identify single nucleotide polymorphisms (SNPs) with minimal technology-specific bias. Leveraging data from over 16,300 ancient individuals, we apply this method to identify a set of a million SNPs that we call the “Compatibility” panel, and which has been effectively assayed in a large fraction of ancient human DNA experiments published to date. We also identify a subset of these SNPs, the “Compatibility-HO” panel, which further restricts to positions that have been assayed in more than ten thousand modern individuals from more than a thousand diverse populations using the Affymetrix Human Origins (HO) genotyping array. The Compatibility panel reduces spurious *Z*-scores due to different sequencing platforms by nearly an order of magnitude, while retaining around 65-85% of statistical power for *f*-statistic analysis. We also provide a tool for users to select different tradeoffs between bias and power as well as sequencing platforms for their specific analyses.

## Introduction

The number of ancient individuals with genome-wide ancient DNA data has expanded exponentially over the past decade and a half (Mallick et al. 2024). These datasets, which crossed the threshold of 10,000 humans with genome-wide data in 2024, have offered granular insights into ancient population history and cultural practices, and have also enabled the detection of natural selection through time-series analysis (Mathieson and Terhorst 2022; Akbari et al. 2024). The increase in sample size has been turbo-charged by in-solution enrichment for targeted subsets of more than a million single nucleotide polymorphisms (SNPs) informative about history and biology, which reduces the required amount of DNA sequencing by one to two orders of magnitude per individual. As of September 2025, the main in-solution SNP enrichment methods used in the field are the “Agilent 1240k reagent” (AG), accounting for about 65% of all published individuals, and the “Twist Ancient DNA” assay (Twist or TW), accounting for 5% of published individuals. While they have greatly decreased the cost of ancient DNA analysis, in-solution enrichment methods have the drawback that they are more prone to biases toward one allele or another at analyzed positions, and yield more variable coverage across targeted SNPs than the approach of shotgun sequencing without an enrichment step (Rohland et al. 2022). These differences have meant that in co-analysis of data produced using different technological platforms, individuals with data produced using the same platform often share alleles at an elevated rate with each other, leading to false-positive inferences of shared ancestry. As a result of these biases, many researchers have restricted many of their analyses to individuals sequenced with the same technology. But since key individuals have been analyzed using different technologies, such restriction significantly limits the scientific questions that can be asked. Datasets from ancient individuals also tend to share alleles at a higher rate than with modern individuals, not due to more shared ancestry, but instead due to the propensity of ancient DNA methods to better capture some alleles than others, making it difficult to co-analyze modern and ancient individuals (Rohland et al. 2022).

Different approaches have been suggested for combining data sequenced using different technologies. A first approach consists of using imputed data (Sousa da Mota et al. 2023; Martiniano et al. 2017). Imputation works by copying haplotype segments from a reference panel, and then selecting the path that best represents the sequencing data. By copying segments from an external panel, this method can leverage neighboring sites to correct for local biases that are due to sequencing. This method has been effectively used to increase the power to discover signals of directional selection in a data set comprising thousands of individuals sequenced with various sequencing platforms (Akbari et al. 2024). However, the approach is restricted to individuals with sufficient sequencing depth (>0.5x genome-wide coverage) and requires the individual’s haplotype variation to be well represented in the reference panel (Sousa da Mota et al. 2023). A second approach focuses on selecting a subset of SNPs that are unbiased across technologies. By selecting SNPs where the ratio of reads mapping to the reference allele at heterozygous sites differed by less than 4% across technologies, one study identified a panel of SNPs comprising 42% of the original 1240k (Rohland et al. 2022), increasing co-analyzability but greatly reducing the number of SNPs and power of downstream analyses. The previously reported SNP panel was also not optimized to remove SNPs that show bias relative to datasets of modern individuals, making co-analysis of modern and ancient individuals with this panel unreliable (Pickrell et al. 2012). Third, SNP loadings in PCA have been used to identify sites that contribute to significant biases in some principal components. This approach requires careful processing to disentangle true ancestral components that are confounded with the sequencing platforms from technical artifacts, but it offers a practical way of performing clustering on a given dataset (Margaryan et al. 2020). Finally, novel tools for pre-processing paleogenomes have been introduced to reduce reference bias. These methods require FASTQ files to be publicly available, which limits their applicability to some legacy datasets (Koptekin et al. 2025).

In this work, we aim to introduce a method that improves upon the previous approach (Rohland et al. 2022). First, we detect biased SNPs using an increased sample size, leveraging over 16,000 individuals with >1x coverage, compared to fewer than 500 for the previously reported co-analysis SNP panel. Second, we use a different statistical test that accounts for the variance in read counts, and does not require the same individuals to be sequenced on different technologies. Third, the statistical test generalizes to an arbitrary number of analysis platforms, and we leverage this to report a “Compatibility” SNP panel that allows unbiased co-analysis of ancient and modern DNA data generated using AG, Twist, low-coverage shotgun (SG), or diploid sequencing platforms. We also identify a subset of these SNPs that are also targeted by the Affymetrix Human Origins (HO) genotyping array, the “Compatibility-HO” panel, and that we verify as producing co-analyzable data between modern and ancient data.

## Results

### Annotating SNPs based on the strength of evidence for incompatibility across platforms

To identify SNPs for the Compatibility panel, we started with autosomal SNPs from the 2M SNP set introduced by Rohland et al. (Rohland et al. 2022), and then removed SNPs based on a series of criteria. First, we removed all SNPs that were not polymorphic in the 1000 Genomes Project (1KGP) (2015) or GnomAD (Chen et al. 2023) datasets. Next, we divided the individuals with sufficient coverage for imputation (c>1x) from our internal database according to their sequencing technology (Table 1). For computational purposes, we only considered the 7,154 individuals with coverage above 2x for the TW technology, which remains the technology with the largest sample size despite this strong minimum coverage threshold. We leveraged haplotypes to identify sites that are likely to be heterozygous (imputation genotype posterior > 0.9), and looked for sites where the read counts did not match this status. To achieve this, we compiled the total read count on each allele for each technology (Supplemental Table S1), and calculated a Bayes Factor to assess if the proportion of reads mapping to each allele is independent of the sequencing technology (see Methods, Supplemental Note). We ran these annotations using all technologies listed in Table 1 for our Default Panel (Supplemental Table S2), or restricting to technologies used to sequence ancient individuals (Supplemental Table S3). Additionally, we annotated the SNPs targeted by the Affymetrix Human Origins array based on their relative mismatch rate between sequencing and genotyping data (Supplemental Table S4, see Methods).

**Table 1.**
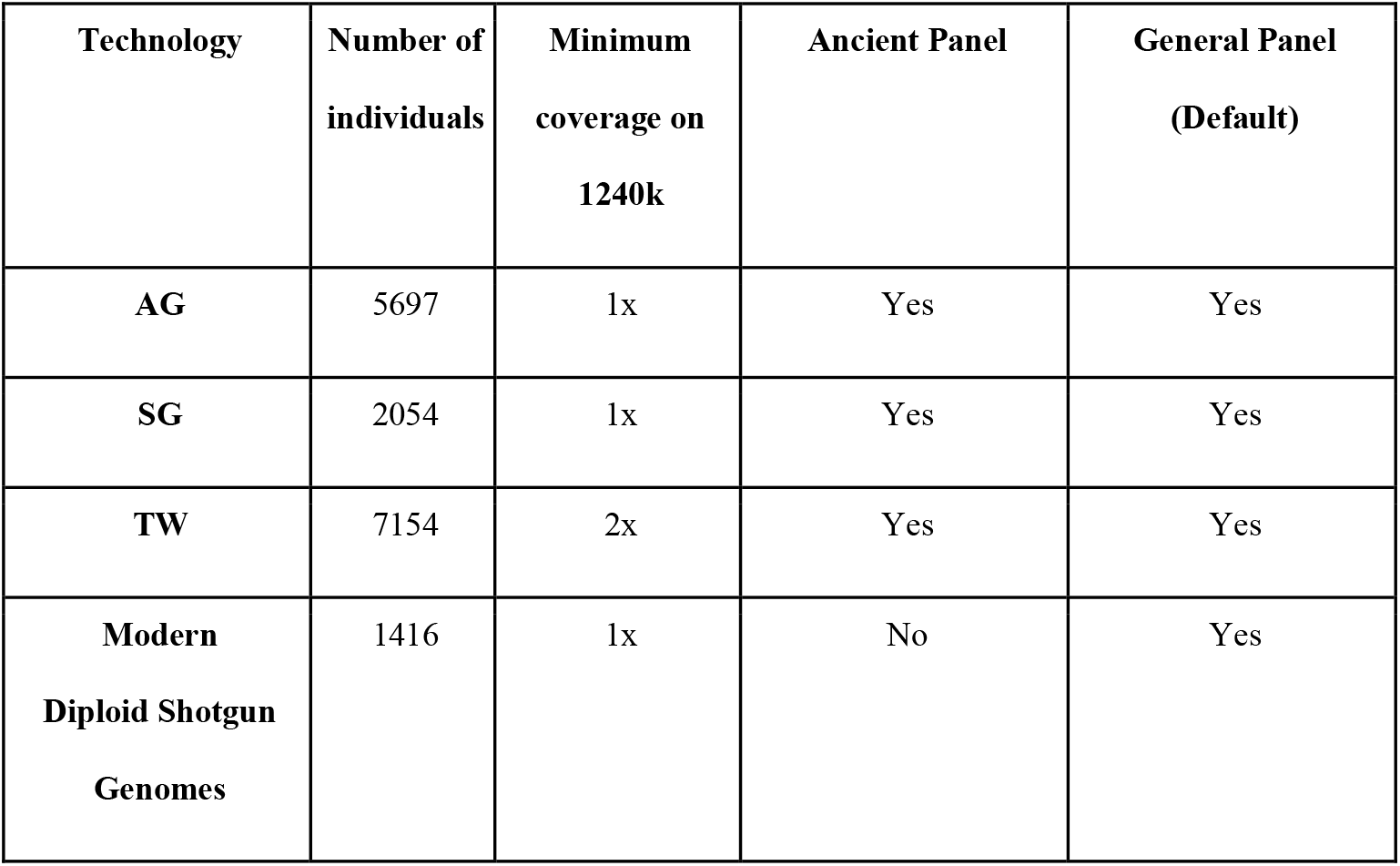
Summary of technologies and sample sizes used in constructing the Compatibility panels. For each technology, we report the number of individuals analyzed and the minimum required coverage for inclusion in the dataset. Summary statistics regarding the number of reads mapping to reference and alternate alleles at each locus are in Supplemental Table S1. The last two columns specify the technologies used for building the Ancient and General Panels.

| Technology | Number of individuals | Minimum coverage on 1240k | Ancient Panel | General Panel (Default) |
| --- | --- | --- | --- | --- |
| AG | 5697 | 1x | Yes | Yes |
| SG | 2054 | 1x | Yes | Yes |
| TW | 7154 | 2x | Yes | Yes |
| Modern Diploid Shotgun Genomes | 1416 | 1x | No | Yes |

We iteratively removed SNPs with the highest bias from the 2M set, and then quantified the remaining bias by computing an *f*_*4*_ statistic on held-out individuals assessed using two different technologies:

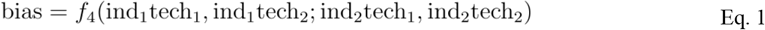

Figure 1a illustrates this remaining bias across different panel sizes. The number of retained SNPs determines the trade-off between statistical power and residual bias in the panel. For the Compatibility panel, we retained SNPs with a Bayes Factor greater than 1, corresponding to SNPs where our approximate model favors the hypothesis of no bias. The remaining bias at various thresholds is depicted in Figure 1a. We also provide a Strict Compatibility panel, retaining only SNPs with a Bayes Factor greater than 3 (Supplemental Figure S1). The same threshold was used to build Ancient-specific Compatibility panels (Table 1). Unless specified otherwise, the Compatibility panel refers to the panel inferred using all technologies.

**Figure 1:**
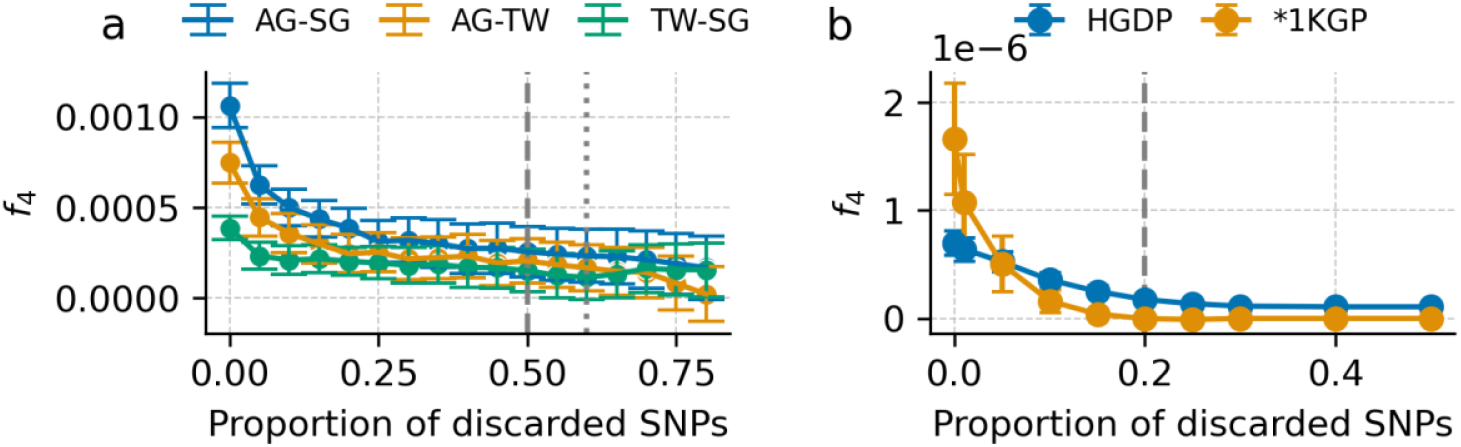
Selection of a threshold for inclusion in the general Compatibility panels. **(a)** Bias, as defined by Eq. 1, for held-out individuals sequenced using at least two technologies among AG, TW and SG. The average value of the f_4_ statistics and 1.96 x s.e.m. are displayed. Vertical dashed and dotted lines indicate thresholds for the Compatibility panel and Strict Compatibility panel, respectively. **(b)** Same quantity for individuals sequenced and genotyped in the Affymetrix Human Origins Array. The individuals from 1KGP were used for the SNP annotation, while the HGDP ones provide true out-of-sample validation.

Next, we used individuals with both shotgun sequencing and HO genotyping data, and annotated each SNP of the HO assay based on the relative mismatch rate between the genotypes obtained from the different technologies (Methods). Using held-out individuals from the HGDP dataset, we calculated the remaining bias using Eq.1. We stopped removing SNPs when the magnitude of the bias reached 0 on the 1KGP individuals used for annotating the SNPs, corresponding to 20% discarded SNPs. At this threshold, there was an 80% reduction of the bias in the held-out HDDP dataset (Figure 1b). The list of SNPs included in the default Compatibility panel is available in Supplemental Table S8, and the various other versions in Supplemental Tables S9-S15.

### Properties of the Compatibility panel

We compared the original and unbiased SNP panels with regard to a series of metrics relevant to population genetic analysis.

We first calculated the fraction of SNPs intersecting the classic 1240k set. For the default Panels, we only observed a mild difference between the original fraction, 0.61, and the new value, 0.60. For the strict panels, however, this proportion rose to 0.67, a significant overrepresentation of these SNPs (Supplemental Table S5, p-value *p<10*^*-5*^). Second, we compared the minor allele frequency (MAF) spectra of both panels using individuals labeled with African (Figure 2a) or European (Figure 2b) ancestry. In both instances, we observed an overrepresentation of SNPs with large MAF (above ∼15%) among the discarded SNPs. This plausibly reflects the fact that fewer individuals are heterozygous at these sites and that the imputation genotype posterior tends to be lower for rare variants, leading to fewer reads available to detect bias for these SNPs. Third and finally, we compared the distribution of mutation profiles, noticing a depletion of transitions in the discarded SNPs (Figure 2c). This might reflect lower imputation genotype posterior probabilities due to postmortem degradation, which would be expected to reduce the number of individuals passing the filtering criteria at these loci. Alternatively, it could reflect systematic differences in the allele frequency spectra of transitions and transversions in the 2M SNP set, for example due to GC-biased gene conversion, or the fact that the ascertainment of SNPs for the Human Origins subcomponent of the 2M SNP panel was non-random with respect to allelic classes that were more or less efficient to design with the genotyping technology(Patterson et al. 2012).

**Figure 2:**
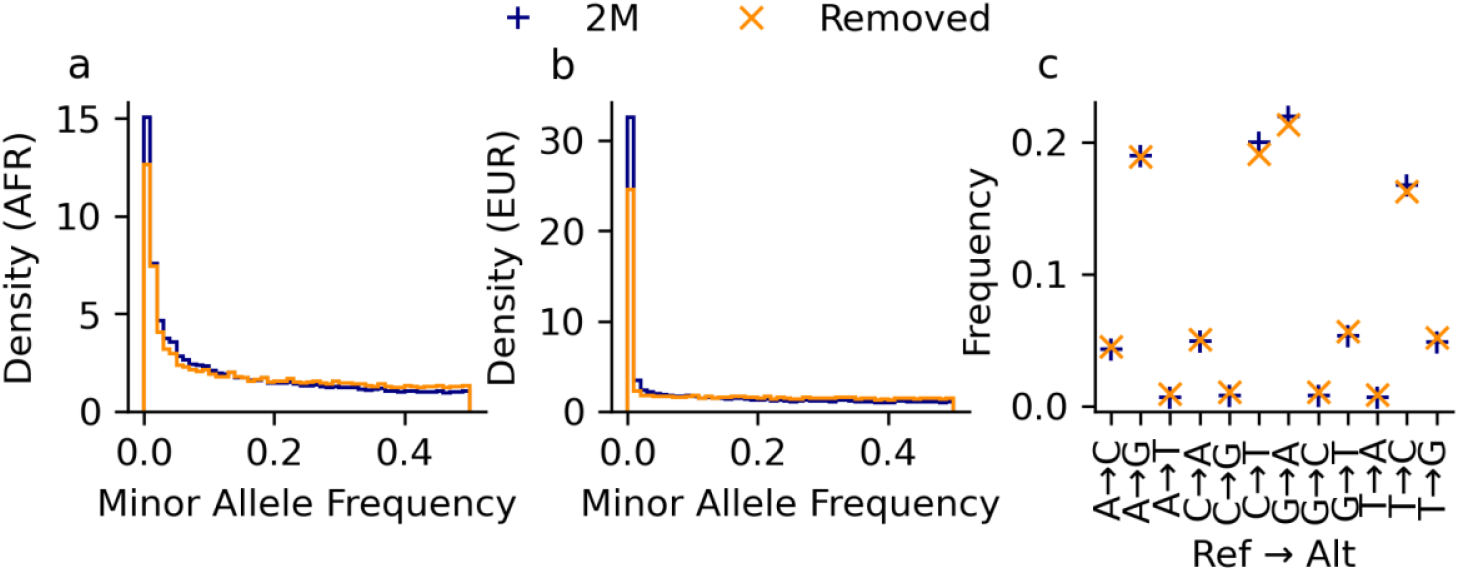
Properties of the Compatibility panel SNPs. **a-b)** Minor allele frequency spectrum in the 1KGP AFR (a) and EUR (b) continental labels. c) Distribution of the mutation profiles in the 2M and Compatibility panels.

### Minimal technical bias in practical settings

We evaluated the bias reduction provided by our SNP panels in realistic scenarios. To achieve this, we calculated three *f*_*4*_(pop1, pop2; pop3, pop4) statistics which are not expected to differ significantly from 0 given the relationships among these groups. In each of the three *f*_*4*_ tests, populations 1 and 3 included only AG data, while populations 2 and 4 consisted of data sequenced using the same alternative technology (Twist, Shotgun, Diploid Modern, or HO). This setup corresponds to a worst-case scenario, where any technology-specific biases can be expected to lead to a particularly significant deviation from 0. Indeed, all *f*_*4*_ tests produced results significantly different from 0 in this setting (*Z-*score*>8*.*7*) (Table 2). Using the recommended SNP panel to compute these *f*_*4*_ statistics showed up to an order of magnitude reduction in the *Z-*scores, with the remaining *Z*-scores reaching values below what is typically reported in such analyses (*Z<3*). These values are even lower when a specific Compatibility panel or the strict versions are used. In these cases, no *Z*-score was nominally significant (Table 2). In real analyses, a more uniform distribution of the sequencing technologies across the populations would decrease the bias. Overall, these results suggest that our proposed SNP panels effectively reduce biases.

**Table 2:** f_4_ statistics obtained using the original and new SNP panels. Data from different technologies were used to calculate f_4_ statistics between different groups. AG-TW: f4(England Neolithic TW (n=16), England Neolithic AG (n=25); Yamnaya TW (n=25), Yamnaya AG (n=25)). AG-SG: f4(England Medieval AG (n=65), England Medieval SG (n=8); Denmark Medieval AG (n=14), Viking SG (n=27)). AG-Modern: f4(YRI(n=101), Kenya Makwasinyi AG(n=10); GBR(n=90), England Medieval AG(n=65)). AG-HO: f4(Kenya Makwasinyi(n=10) AG, Kikuyu HO(n=4); Belgium Medieval AG(n=36), French HO(n=61)). We used the default and strict compatibility panels to calculate the Z-score (General), together with panels specifically built for each pair of technologies (the Ancient Panel for the first two rows, and the AG-Modern for the last two). The last row shows data further restricted to the HO compatible sites, as described in the main text.

| Tech 1 | Tech 2 | N(tot) | Z(2M) | Z(General) |  | Z(Specific) |  |
| --- | --- | --- | --- | --- | --- | --- | --- |
|  |  |  |  | Default | Strict | Default | Strict |
| AG | TW | 91 | 10.8 | 1.7 | 1.41 | 1.7 | 1.06 |
| AG | SG | 114 | 12.2 | 1 | 0.57 | 0.8 | 0.51 |
| AG | Modern | 266 | 11.3 | 2.39 | 1.91 | 1.73 | 1.55 |
| AG | HO | 111 | 8.7 | 0.73 | 0.39 | 0.06 | -0.12 |

Finally, we assessed how the compatibility panel affects analyses beyond *f*_*4*_ statistics. We projected 159 ancient individuals with both AG and SG data onto principal components computed using modern West Eurasian populations (Supplemental Table S6, Methods). We then calculated the mean difference between the two projections of each individual for each of the PCs (Supplemental Figure S2). When the original HO SNP set was used, we observed significant biases in PCs 1, 3, 4, and 5 (*p* = 4.4×10^-4^, 1.9×10^-2^, 2.8×10^-5^, and 8.5×10^-3^, respectively). The bias was no longer significant using the compatibility SNP set for these PCs, although a small residual artifact persisted on PC 10, which remained nominally significant (*p* = 3.8×10^-2^).

### Minimal reduction in statistical power from use of the Compatibility panel

We quantified the reduction in statistical power resulting from the lower number of SNPs in the Compatibility panel. To perform this computation, we re-examined a dataset of 291 individuals assigned to different groups across Iron Age Britain (Patterson et al. 2022). Following the original study, we first tested for differences in ancestry between pairs of groups by calculating statistics of the form *f*_*4*_(group 1, group 2; Turkey Neolithic, Old Steppe), restricting to the pairs of groups with Z-scores above 3 in the original study. We calculated this quantity using both the original 1240k panel and our Compatibility panels. We then fit a regression line between these points, and observed a regression coefficient of 0.82 for the Compatibility panel, and 0.86 for the Ancient-specific one (Figure 3a). The same analysis performed for the restricted version of these panels led to regression coefficients of 0.83 and 0.78, respectively (Supplemental Figure S2).

**Figure 3:**
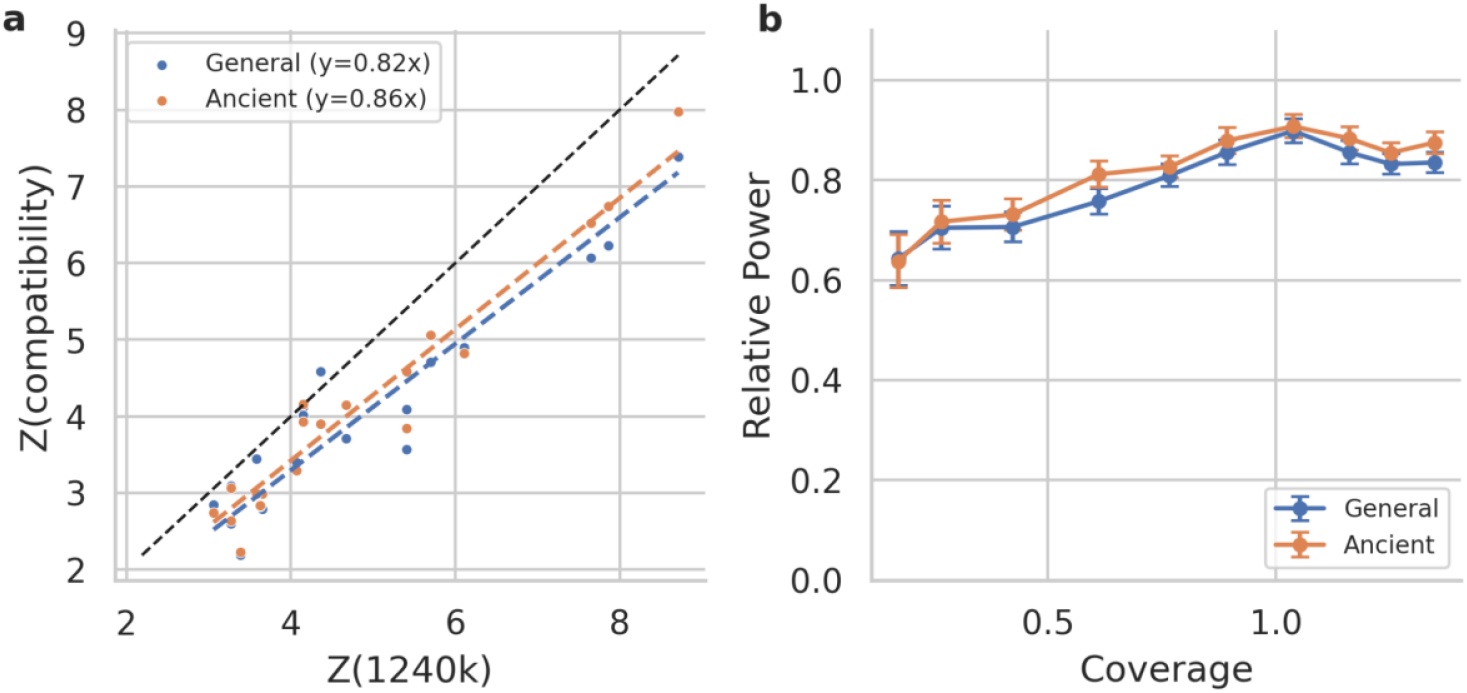
Power reduction in the new SNP set. **(a)** Differences in Z-scores obtained using the original SNP set (1240k) and the Compatibility panels. The regression lines y=0.82x and y=0.86x are shown to illustrate the reduction in power of the General and Ancient Compatibility panels, respectively. **(b)** Same analysis, but stratifying the data by coverage. The relative power corresponds to the regression coefficient fitted on Z(Compatibility) = a Z(1240k). Error bars correspond to the 95% CI around the parameter a, assuming independent Z-scores (Methods).

We analyzed statistics of the same form, but replacing British Iron Age groups with British Iron Age individuals, to assess how coverage impacts the standard errors of the *f*_*4*_ statistics. To do so, we calculated *f*_*4*_(ind 1, ind 2; Turkey Neolithic, Old Steppe) for pairs of individuals belonging to pairs of groups that were detected as being significantly different in their ancestry in the original study (reported absolute *Z-*scores above 3). For each pair, we recorded the number of overlapping SNPs, as well as the *Z-*scores obtained when calculating the *f*_*4*_ statistics with the new and original panels. We then split all results according to the number of overlapping sites, creating 10 sets of the same size. Within each group, we estimated the regression coefficient of the *Z-*scores calculated on both panels, as we did in the previous analysis (see Methods). The regression coefficient (Figure 3b) stays at 0.85 for individuals with coverage > 0.8× before progressively dropping to 0.65. The greater loss of power for lower coverage individuals reflects the fact that for such individuals, there is expected to be little linkage disequilibrium among SNPs for which data are available, and hence nearby SNPs are not filling in information gaps as is the case for higher coverage individuals.

## Discussion

We released Compatibility panels, which are a subset of a previously described “2M” panel consisting of the union of positions targeted by the AG and Twist assays, along with SNPs that were effectively captured using either of these assays due to being within 50 base pairs of targeted positions, and 15 multiallelic SNPs and 6 indels (Rohland et al. 2022). The default Compatibility panel comprises 913,708 autosomal SNPs, to which we added the unfiltered 63,994 positions on the X chromosome and 199,881 positions on the Y chromosome from the 2M panel.

By reducing biases across technologies, our curated SNP panels can be used for multiple downstream analyses. One application is preserving compatibility with data generated using earlier sequencing technologies. As a result, new individuals can be analyzed without needing to match the platforms used in previous studies. Second, the panels enable joint modeling of ancient and modern individuals in -based methods, such as qpAdm and qpWave(Haak et al. 2015). Finally, methods beyond statistics may benefit from the curated panels. This is particularly relevant for frequency-based approaches and principal component analyses, where platform-specific biases can lead to spurious results.

A limitation of our model is that reads are treated as independent entities. This prevents strict probabilistic interpretation of the annotations, and requires relying on external validation sets to test the panels, as done in this analysis. As more individuals become publicly available, allowing for the use of individual read counts, a mixed model accounting for individual-level effects could increase the power of the compatibility panels. Next, the power to detect technology-specific bias is directly linked to the number of heterozygous individuals sequenced at each locus. As a result, our method is underpowered to detect problematic SNPs that have minor allele frequencies under 10%, or that are only present in underrepresented ancestries. An important caveat is that the new SNP panel, despite significantly reducing bias, does not fully eliminate it. Therefore, we recommend that analysts be cautious when interpreting *Z-*scores that are close to the rejection threshold. Finally, the compatibility panel has non-homogeneous coverage across technologies. For some analyses, it might be valuable to impose a further coverage threshold on the remaining SNPs.

As larger and more diverse datasets become available, future updates of this panel could further reduce biases. We also release a script and current read counts that researchers can use to build their own compatibility panel. For example, if residual inflation in *Z-*scores is observed, analysts can further reduce the number of SNPs in the compatibility panel, create a panel restricted to the technologies present in their analysis, or generate a population-specific panel by only using reads from individuals in relevant populations. For convenience, the package contains the lists obtained for common pairs of technologies and various thresholds, in addition to the recommended ones.

Finally, for some analyses, the panels we provide will not be useful. Whole genome sequencing remains critical for many applications such as de novo SNP discovery, methylation studies and structural variant detection.

## Methods

### Removing SNPs based on differences in reference biases

Our analysis leverages more than 16,300 individuals (Table 1), who were imputed using the same pipeline as that described in a recent study of natural selection (Akbari et al. 2024). For each SNP in the 2M SNP panel (Rohland et al. 2022) that is polymorphic in the 1KGP or Gnomad datasets (using the GRCh37 human reference genome), we retained individuals imputed as heterozygous with a genotype posterior probability above 90%. Using these individuals, we reported, for each technology, the number of reads mapping to the reference and alternate alleles (Supplemental Table 1). Because most of the individuals used for this study are unpublished, we could not report individual-level read counts, and our model thus considers each read as an independent entity. We then built a contingency table with sequencing technologies as rows and the number of reads mapping to the reference and alternate alleles as columns. For each SNP, we calculated a Bayes Factor assessing the independence of each row (Supplemental Note). We decided to provide panels for two distinct thresholds, a Bayes Factor of 1 and 3. While these numbers are commonly used in the literature and worked well in our tests, users can apply different thresholds using our Python package.

### Removing SNPs for the Compatibility-HO panel

We leveraged 1028 individuals with both sequencing and genotyping data(Lazaridis et al. 2014; Mallick et al. 2016; Skoglund et al. 2016; Prüfer et al. 2014; Fan et al. 2019; 2015; Patterson et al. 2012; Bergström et al. 2020; Nakatsuka et al. 2017). For each SNP, we calculated the relative number of mismatches over the total number of heterozygous individuals at that site. We discarded SNPs according to this metric. The threshold was set to the first value for which no bias was observed in the 1KGP individuals.

### Calculating *f*_*4*_ statistics

We calculated *f*_*4*_ statistics using Admixtools2 (Maier et al. 2023) with the ‘maxmiss’ parameter set to 0.

### PCA

We ran a PCA using SmartPCA(Patterson et al. 2006) with the lsqproject option set to YES. We calculated principal components using the individuals of Supplemental Table S6 with the HO, and then projected the ancient individual on these axes. We then used a t-test to calculate p-values for deviation from a zero-mean difference on each component.

### Power Analysis

We analyzed the same set of individuals and their corresponding population labels as in Patterson et al. 2022(Patterson et al. 2022)(Supplemental Table S7). We fit a linear regression without an intercept to the resulting *Z-*scores to estimate the power reduction. For the coverage analysis, we proceeded in the same way but this time using pairs of individuals belonging to the different groups with large Z-scores (>3) . We separated the pairs of individuals according to the number of overlapping sites. If *p* denotes the proportion of overlapping sites on the 1240k panel, and *c* denote the coverage, we can use a Poisson approximation to obtain:

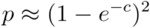

We inverted this relation to obtain *c*. This equation assumes that the probability of a SNP being missing is independent of other reads and uniform, which means that the actual value of *c* is likely to be lower. In other words, the power is likely to remain higher at lower coverages than the reported 0.8x.

We fitted the regression line assuming all *Z-*scores are independent. This means that the confidence intervals provided around the slope are tighter than what is expected if points were truly independent.

### Editing

We wrote a rough draft of each paragraph, and once completed, used Large Language Models to edit at the sentence level. We typically asked for three versions of some sentences. We then selected the version we preferred, or we mixed different elements of different answers. This tool was only used to edit the style of this manuscript, not its structure nor its content. We then carried out multiple additional rounds of editing (not assisted by these tools).

## Code Availability

We released a Python package (https://github.com/rmnfournier/compatibility-panel) for users to create their own panels for specific use cases.

## Data Availability

Read counts for each technology are available in Supplemental Table S1. Annotations for the default and ancient-specific panels are available in Supplemental Tables S2 and S3, respectively. Compatibility panels used in the manuscript are available in Supplemental Tables S8 (default panel) to S15. The list of individuals used for the analyses is available in Supplemental Tables S6 and S7, and can be accessed through the AADR dataset (Mallick et al. 2024).

## Acknowledgments

We thank Shop Mallick, Bárbara Sousa da Mota, and Nick Patterson for valuable discussions. We thank reviewers for their valuable insights. We thank the bioinformatics team in our laboratory for making available the datasets we used for identifying SNPs susceptible to bias. This work was supported by John Templeton Foundation grant 61220; by National Institutes of Health grant HG012287; and by the Howard Hughes Medical Institute.

## Notes

### Competing Interest Statement

The authors have declared no competing interest.

### Summary of Updates

Improved summary statistics for low MAF variants and revisions

