## Supplementary figures and images for "A SNP panel for co-analysis of capture and shotgun ancient DNA data"

### Supplemental Figure S1

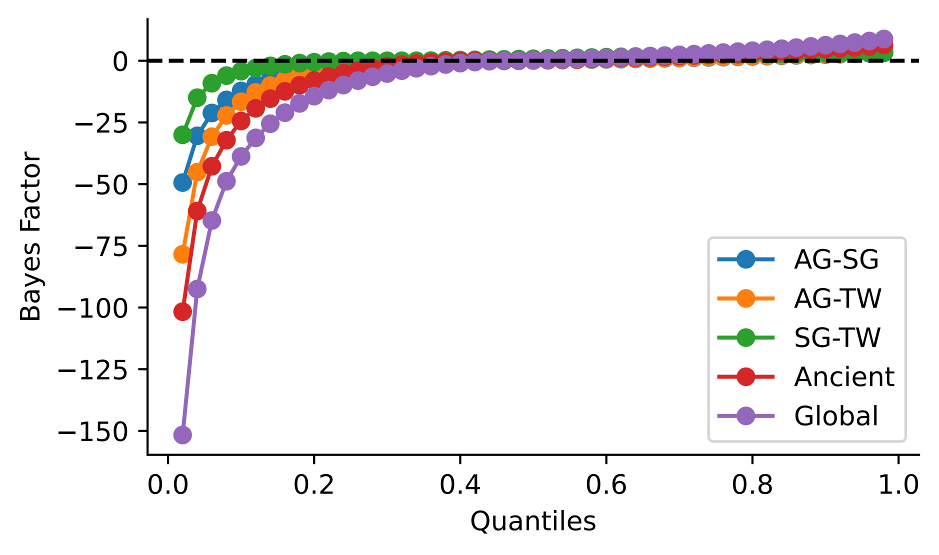

### Supplemental Figure S2

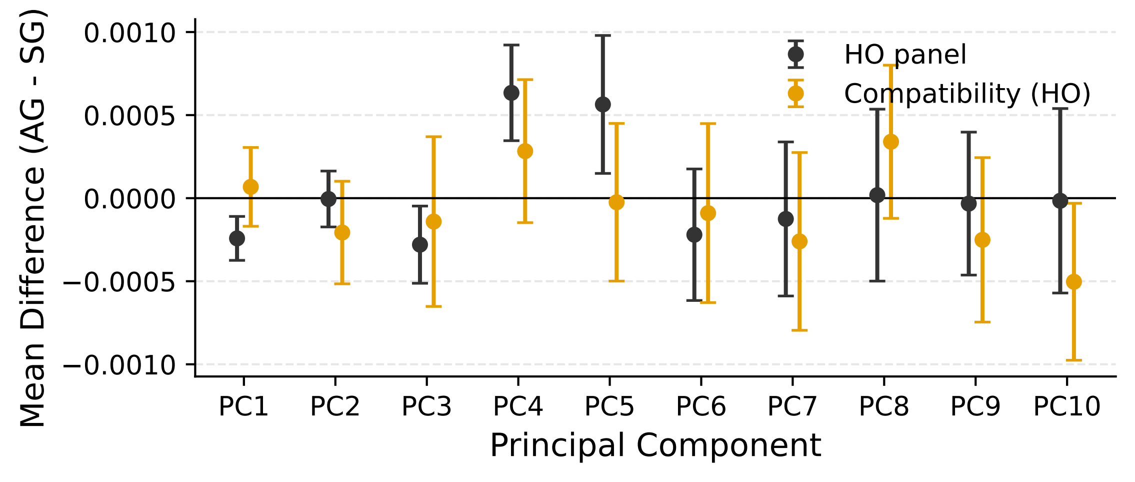

### Supplemental Figure S3

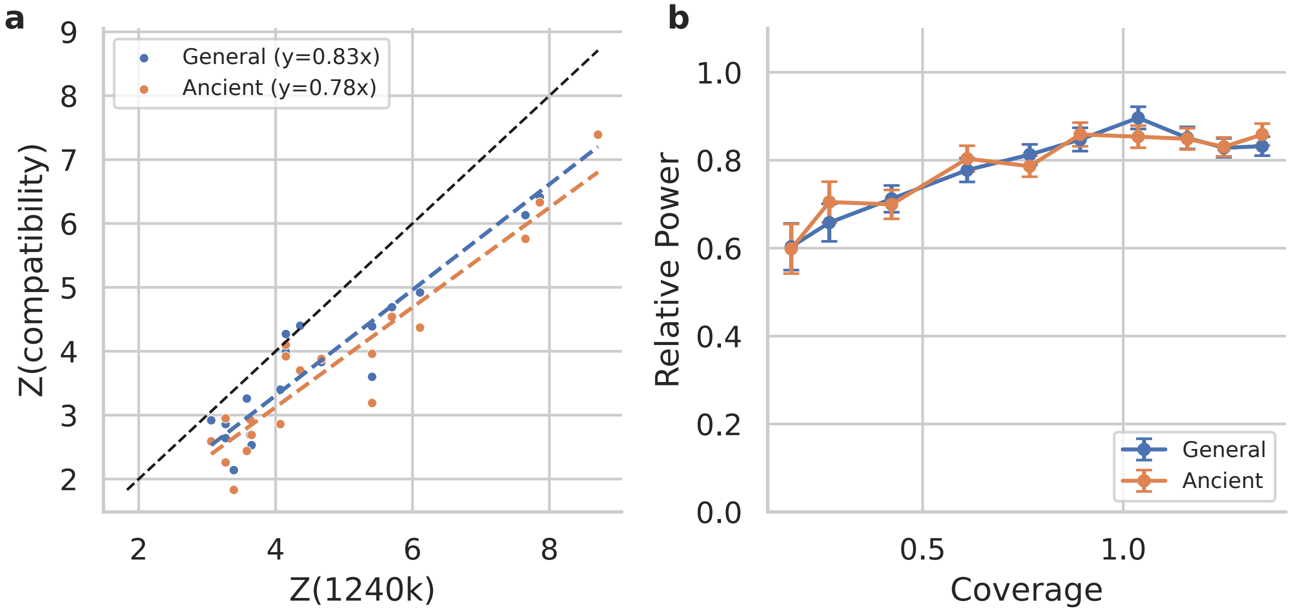
