## Supplemental Note for "A SNP panel for co-analysis of capture and shotgun ancient DNA data"

### Compatibility Panel: Supplementary Note

Romain Fournier

December 2025

#### 1 Annotations

For a given SNP, let $\mathbf{n} = \{n_1, n_2, \dots\}$ be the total number of reads for each technology, and $\mathbf{k} = \{k_1, k_2, \dots\}$ be the number of reads mapping to the reference allele among those. As a reminder, only heterozygous individuals are considered for this analysis. We calculate the likelihood of observing these counts under two hypotheses. For the first model, $H_0$, we assume that the proportion of reads mapping to the reference allele is the same across technologies, albeit unknown. For the second model, $H_1$, we assume that each technology has its own ratio.

Let $K_{total} = \sum_{i=1}^{t} k_i$ and $N_{total} = \sum_{i=1}^{t} n_i$. Assuming known mapping probability $p$, the likelihood under $H_0$ is given by:

$$P(\mathbf{k}|\mathbf{n}, p) = \prod_{i=1}^{t} \left[ \binom{n_i}{k_i} p^{k_i} (1-p)^{n_i - k_i} \right] = \left[ \prod_{i=1}^{t} \binom{n_i}{k_i} \right] p^{K_{total}} (1-p)^{N_{total} - K_{total}}$$

Assuming a $B(2,2)$ (beta) prior, favoring the expected 50-50 ratio, the model evidence for $H_0$ is given by:

$$P(\mathbf{k}|\mathbf{n}, H_0) = \left[ \prod_{i=1}^{t} \binom{n_i}{k_i} \right] \frac{B(K_{total} + 2, N_{total} - K_{total} + 2)}{B(2,2)}$$

For $H_1$, we treat each row as having an independent mapping ratio. The model evidence is thus:

$$P(\mathbf{k}|\mathbf{n}, H_1) = \prod_{i=1}^{t} \left[ \binom{n_i}{k_i} \frac{B(k_i + 2, n_i - k_i + 2)}{B(2,2)} \right]$$

Finally, we calculate the Bayes Factor, defined as the ratio of the model evidences:

$$BF = \frac{B(K_{total} + 2, N_{total} - K_{total} + 2)}{B(2,2)} \times \left[ \prod_{i=1}^{t} \frac{B(k_i + 2, n_i - k_i + 2)}{B(2,2)} \right]^{-1} \quad (1)$$

.

#### 2 Supplementary Figures

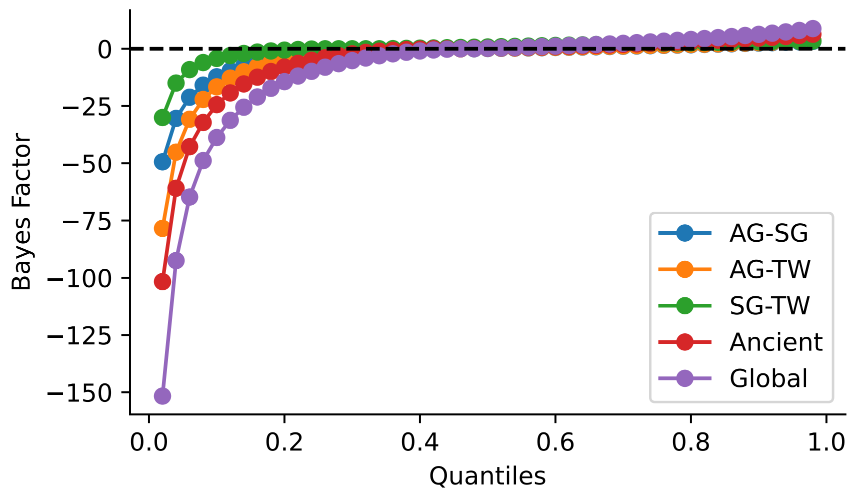


Supplementary Figure S1: Quantiles of the log Bayes Factor (Eq. 1) calculated for different panels. The Ancient Panel contains reads from individuals sequenced with AG, SG, and TW technologies, and the General Panel adds reads from modern individuals (Table 1).

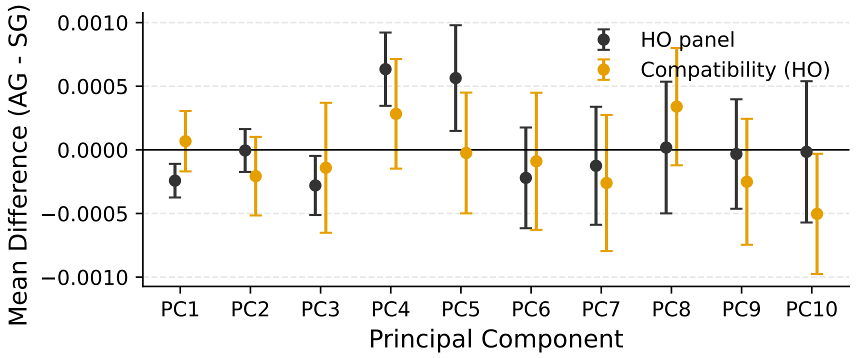


Supplementary Figure S2: Individuals with both AG and SG were projected onto principal components calculated using modern HO individuals. The mean differences for each coordinate, along with 95% CIs (1.96 × s.e.m.), are shown for projections using SNPs from the original HO panel and restricted to the default Compatibility Panel.

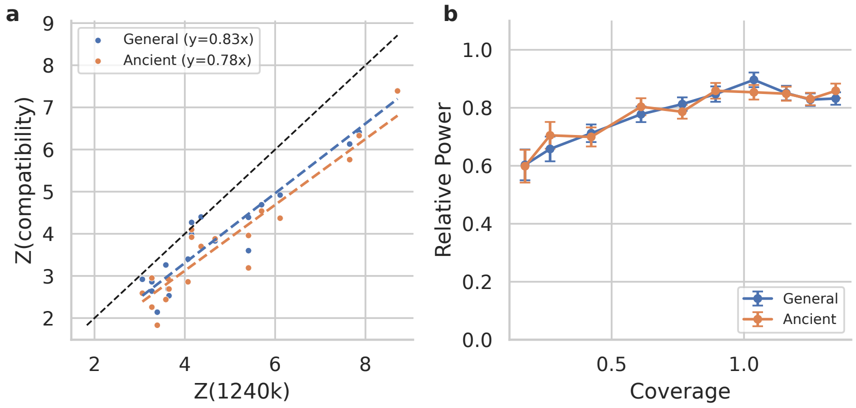


Supplementary Figure S3: Power reduction in the Strict SNP sets. (a) Differences in $Z$-scores obtained using the original SNP set (1240k) and the Strict Compatibility panels. The regression lines $y = 0.83x$ and $y = 0.78x$ are shown to illustrate the reduction in power of the General and Ancient Compatibility panels, respectively. (b) Same analysis, but stratifying the data by coverage. The relative power corresponds to the regression coefficient fitted on $Z(\text{Compatibility}) = \text{a}\ Z(\text{1240k})$. Error bars correspond to the 95% CI around the parameter a, assuming independent $Z$-scores (Methods).
